# IDENTIFICATION AND CHARACTERIZATION OF A LARGE SOURCE OF PRIMARY MESENCHYMAL STEM CELLS TIGHTLY ADHERED TO BONE SURFACES OF HUMAN VERTEBRAL BODY MARROW CAVITIES

**DOI:** 10.1101/2020.05.04.076950

**Authors:** Brian H. Johnstone, Hannah M. Miller, Madelyn R. Beck, Dongsheng Gu, Sreedhar Thirumala, Michael LaFontaine, Gerald Brandacher, Erik J. Woods

## Abstract

Therapeutic allogeneic mesenchymal stem/stromal cells (MSC) are currently in clinical trials for evaluating their effectiveness in treating many different disease indications. Eventual commercialization for broad distribution will require further improvements in manufacturing processes to economically manufacture MSC at sufficient scales required to satisfy projected demands. A key contributor to the present high cost of goods (COG) for MSC manufacturing is the need to create master cell banks (MCBs) from multiple donors, which leads to variability in large-scale manufacturing runs. Therefore, the availability of large single donor depots of primary MSC would greatly benefit the cell therapy market by reducing costs associated with manufacturing.

We have discovered that an abundant population of cells possessing all the hallmarks of MSC is tightly associated with the vertebral body (VB) bone matrix and are only liberated by proteolytic digestion. Here we demonstrate that these vertebral bone-adherent (vBA) MSC possess all the International Society of Cell and Gene Therapy (ISCT)-defined characteristics (e.g., plastic adherence, surface marker expression, and trilineage differentiation) of MSC and, therefore, have termed them vBA-MSC, to distinguish this population from loosely associated MSC recovered through aspiration or rinsing of the bone marrow (BM) compartment.

Pilot banking and expansion was performed with vBA-MSC obtained from 3 deceased donors and it was demonstrated that bank sizes averaging 2.9×10^8^ ± 1.35×10^8^ vBA-MSC at passage one were obtainable from only 5 g of digested VB bone fragments. Each bank of cells demonstrated robust proliferation through a total of 9 passages without significant reduction in population doubling times. The theoretical total cell yield from the entire amount of bone fragments (approximately 300g) from each donor with limited expansion through 4 passages is 100 trillion (1×10^14^) vBA-MSC, equating to over 10^5^ doses at 10×10^6^ cells/kg for an average 70 kg recipient. Thus, we have established a novel and plentiful source of MSC which will benefit the cell therapy market by overcoming manufacturing and regulatory inefficiencies due to donor-to-donor variability.

## INTRODUCTION

The potent activity as well as high expandability of mesenchymal stem/stromal cells (MSC) has generated considerable interest from commercial entities in developing “off-the-shelf” allogeneic MSC therapeutics derived from a limited number of donors. Development of a cellular therapy based on allogeneic “universal donors” allows for controlled manufacture with careful attention to thoroughly assessing quality (e.g., identity, potency, purity, and safety) of each manufactured lot at significant cost savings compared to manufacturing individual lots of autologous cells from individual donors, such as currently occurs for chimeric antigen receptor (CAR) T cell therapies.

The challenges inherent to manufacturing cellular therapies scale with the size of a manufacturing run. Effective doses of MSC for some indications are as high as 1×10^9^ cells per dose, which would require manufacturing 10 trillion (10×10^12^) cells per year to affordably meet potential demand [1–4]. Even at this level of production, with presumed economies of scale, the cost of goods sold (COGS) per dose of MSC could exceed $100,000 [3]. A significant driver of manufacturing costs, which is amplified proportionately with lot size, is the need to replenish master cell banks (MCB) through isolation of MSC from new donors due to the limited volumes of tissues and fluids that can be safely obtained from healthy volunteers and the limited expansion potential of MSC isolated from each donor [5, 6]. MSC are rare in all tissues, comprising, for instance, ~0.001-0.01% of total nucleated cells (TNC) in BM aspirates [7]. Given that BM aspirates from healthy volunteers are limited for the safety of the donor to 100 ml (50 ml bilaterally from iliac crests), the total yield of fresh, non-passaged MSC is approximately 2×10^4^/donor. Expansion to a trillion cells would require seed stocks of 1×10^7^ MSC in order to limit cell proliferation to 9 population doublings [8]. This number is in addition to the cells reserved for quality control measurements of the expanded MCB and working cell bank (WCB). Thus, the number of MSC obtainable from each donor is more than 3 orders of magnitude less than is optimal for the initial stages of expansion.

The need to constantly replenish cell banks by obtaining fresh cells from new donors introduces inconsistencies into the manufacturing process due to the observed variability between MSC derived from different donors otherwise matched for attributes such as age and health status [6, 9, 10]. Donor-to-donor variability and the resulting economic impact on manufacturing costs is substantial. In one study that examined large scale manufacturing of multiple lots of MSC derived from different donors, it was found that cumulative population doublings between 5 different BM donors varied by 1.8-fold during 30 days in culture [9]. The result was a >13 day variation in process time to manufacture a batch of 350 million MSC. Besides the logistical burden to coordinate batch runs, there was a commensurate increased cost of growth medium, which is also a key cost-driver for cell-based therapy manufacturing [1, 3, 8]. Furthermore, the authors found that there was >18% difference in colony forming potential and >50% difference in interleukin 6 expression, adding an additional complication to quality control verification of potency for each batch derived from individual donors. Similarly, a single center experience with clinical manufacture of 68 batches of MSC from BM recovered from 59 human volunteer donors observed population doubling times that varied by over 2-fold (46.8-141 hours), averaging 71.7 hours, yielding final batch numbers of MSC ranging from 1.9×10^7^ to 5.43×10^9^ (average 5.46×10^8^) [10].

Besides imposing a direct economic burden of increasing COG per manufacturing run, there is also a regulatory burden with associated costs resulting from the need to refresh cell banks. The MCB serves as the reserve of starter cultures for all manufacturing runs using cells from a particular donor. The regulatory requirements for quality and safety assessments of the MCB are costly and time consuming [11]. Of the three overarching parameters (i.e., safety, potency and identity) required to assess suitability of a manufactured cell therapy product, potency as it relates to individual donor characteristics, is most problematic due to the changing profile that occurs with expansion, as described above. This is particularly the case as MSC populations near the limits of expansion and enter into senescence which severely limits their potency [12]. For these reasons, population doubling limit is an important factor for regulatory authorities; albeit, one that is not commonly addressed in filings with FDA [13].

Reducing the economic and regulatory burden of generating multiple MCB lots annually to fulfill the need for large scale manufacturing requires identifying large depots of unmanipulated MSC. Potential solutions could come from abundant tissues harboring MSC that are normally discarded following routine medical procedures or are obtainable post-mortem. Adipose-derived stem/stromal cells (ASC) are obtained from elective procedures that commonly yields liters of tissue and have recently been extensively investigated; however, primarily for autologous uses [14, 15]. Isolation directly from medullary cavity-containing bones obtained through medical procedures or cadavers yields higher percentages of MSC (~0.04%) than are present in aspirates, most likely reflecting lack of peripheral blood contamination [16]. Total nucleated cell counts of ~5×10^9^ have been obtained from BM of vertebral bodies (VB) recovered from deceased organ donors, with each VB containing ~2×10^6^ MSC, or ~2×10^7^ total MSC per typical spinal 9 VB segment recovered [17]. In addition, the ilia, sternum, ribs and heads of long bones are sources of BM from which MSC can be recovered [18–20]. Thus, the VB compartment of BM alone from a typical deceased donor yields >3 orders of magnitude more MSC than can be obtained from a healthy human donor.

In addition to cells obtained by eluting or aspirating BM, another population of MSC is tightly associated with medullary cavity bone structures [21–23]. First identified in rodent long bones, bone-adherent MSC (BA-MSC) have subsequently been isolated from human bone fragments obtained from long bone condyles and vertebrae [24]. We have discovered another source of MSC, termed vertebral BA-MSC (vBA-MSC), which remain attached to fragments of VB bone after extensive washing to remove BM cells and can only be liberated by digestion with proteases. The frequency and functionality of vBA-MSC is equivalent to that in eluted VB BM-MSC. Here we present these data and establish a new source of MSC that could be used in large scale manufacturing processes to produce batches totaling of over a quadrillion cells from an individual donor; thus, satisfying the most optimistic levels of demand for decades and overcoming a current impediment to commercial scale production [2, 8].

## MATERIALS AND METHODS

### Sources of tissues and cells

Vertebrae were recovered from brain-dead organ donors after obtaining informed consent for research use. Each recovered vertebra was assigned a unique identifier. The inclusion criteria for donor selection were brain death, age between 12 and 55 years, non-septicemic, and disease and pathogen free. Live donor aspirated BM from three healthy volunteers was purchased from Lonza (Walkersville, MD). Expanded live donor MSC, cryopreserved at passage 2, were purchased from Lonza. Relevant donor characteristics are presented in Table 1.

**Table 1.** Description of donors used in this study

| **Donor ID** | **Recovered MSC** | **Age** | **Sex** | **Race/ Ethnicity** |
| --- | --- | --- | --- | --- |
| DD1<sup>1</sup> | vBA-MSC, vBM-MSC | 22 | M | Caucasian |
| DD2 | vBA-MSC, vBM-MSC | 13 | M | Caucasian |
| DD3 | vBA-MSC | 35 | M | Hispanic |
| DD4 | vBA-MSC, vBM-MSC | 19 | M | Hispanic |
| DD5 | vBA-MSC | 17 | M | Caucasian |
| DD6 | vBA-MSC | 14 | M | Caucasian |
| DD7 | vBA-MSC | 23 | M | Caucasian |
| LD1 | LD BM-MSC | 20 | F | African American |
| LD2 | LD BM-MSC | 23 | F | African American |
| LD3 | LD BM-MSC | 28 | M | African American |
| LD4 | Ex LD BM-MSC | 24 | F | African American |
| LD5 | Ex LD BM-MSC | 36 | M | African American |
| LD6 | Ex LD BM-MSC | 25 | M | African American |

<sup>1</sup>Abbreviations: DD, deceased donor; LD, live donor; vBM-MSC, vertebral bone marrow-MSC; vBA-MSC, vertebral bone-adherent MSC,

### Deceased Donor Tissue Procurement and Transport

Previously developed clinical recovery methods [16, 25] combined with subsequent experience in the ongoing VCA transplant immunomodulation clinical trial (ClinicalTrials.gov Identifier: NCT01459107) at Johns Hopkins University formed the basis for the procurement and transport protocols. A streamlined organ procurement agency (OPO) recovery procedure, combined with dedicated kits and centralized training on recovery and shipment procedures were employed. Recovered bones were shipped to Ossium Health (Indianapolis, IN). Vertebral sections were procured by six different OPO partners: Gift of Hope (Itasca, IL); Donor Alliance (Denver, CO); Iowa Donor Network (North Liberty, IA); Mid America Transplant (St. Louis, MO); and Nevada Donor Network (Las Vegas, NV). Bones were recovered by OPO personnel using an osteotome and mallet under an IRB approved protocol from research-consented organ and tissue donors. Recovered bones were wrapped in lap sponges and towels soaked in saline to ensure moisture retention during shipment. Wrapped specimens were shipped overnight on wet ice to Ossium's processing facility.

### Manual Debriding

Upon receipt, in a Biological Safety Cabinet, soft tissue was manually debrided using scalpels and gouges. Once visible, the pedicles were removed using a Stryker System 6 Saw (Stryker, Kalamazoo, MI) leaving only the connected vertebral bodies. Vertebral bodies were separated, and intervertebral disc and soft tissue were removed with a scalpel. Care was taken to ensure that the cortical bone was not breached to preserve and protect the hypoxic cancellous bone marrow throughout the entire debriding process.

Using custom-made surgical grade stainless-steel anvil shears, VBs were cut into approximately 5 cm^3^ pieces small enough to be fragmented with a bone grinder. The pieces were immediately submerged into 500 mL processing medium comprised of Plasma-Lyte A pH 7.4 (Baxter Healthcare, Deerfield, IL), containing 2.5% human serum albumin (HSA; Octapharma USA Inc., Hoboken, NJ), 3 U/ml Benzonase (EMD Millipore, Burlington, MA), and 10 U/ml heparin (McKesson, Irving, TX).

### Grinding and Elution

A bone grinder (Biorep Technologies. Inc, Miami, FL) was assembled in a biological safety cabinet. A two-liter stainless steel beaker containing approximately 250 mL of fresh processing medium was placed under the grinding head to catch bone chips and media flow-through. A stainless steel plunger was used to aid in pushing pieces through the grinder. Rinsing through the grinder with processing medium prevented bone pieces from drying out and sticking to the chamber. Once all bone pieces were ground, the chamber was thoroughly rinsed with fresh processing medium. The final volume in the stainless-steel beaker was one liter.

Filtering was performed using bone marrow collection kits with flexible pre-filter and inline filters (Fresenius Kabi, Lake Zurich, IL). All bone grindings and media were carefully transferred to the bone marrow collection kit. The grindings were gently massaged to allow for optimal cell release from grindings. The media was then filtered using two 500 μm and two 200 μm filters. The bone grindings are rinsed using two 500 mL washings with rinse media. Rinse medium was Plasma-Lyte with 2.5% HSA. All bone marrow was then collected in a collection bag. The bone fragments remaining in the filtration kit were stored at 4°C overnight before processing to recover vBA-MSC.

### Digestion protocol for vBA-MSC isolation

Bone fragments (either 1 or 5g) were transferred to sterile 50 ml conical centrifuge tubes. A solution of DE10 collagenase (2 mg/ml; Vitacyte, Indianapolis, IN) was added to the bone fragments at a ratio of 5:1 (volume:weight). The tubes were transferred to a shaking incubator and incubated for 1.5 hrs at 37°C while shaking at 125 rpm. Protease activity was neutralized by adding 2% Stemulate (Cook Regentec, Indianapolis, IN) and suspensions were filtered through a 70 μm cap filter into 50 ml conical screw cap tubes. The filter-retained bone fragments were washed with 25 ml Dulbecco's modified phosphate buffered saline (DPBS) solution containing heparin (10U/ml) and Benzonase (100 U/ml; Millipore, Burlington, MA) which was combined with the original filtrate. Tubes were centrifuged at 350xg for 5 minutes, supernatant aspirated, and the pellets were resuspended in 25 ml DPBS. The suspension was centrifuged again at 350xg for 5 minutes, the supernatant was aspirated, and the pellet was resuspended in 10 ml DPBS for analysis.

### Isolation of MSC from iliac and VB BM

An 1 ml aliquot of concentrated, eluted BM was removed and pipetted into a 50 ml conical vial along with 49 ml DPBS. The vial was centrifuged at 300xg for 10 minutes, supernatant aspirated, and the pellet resuspended in 10 ml Rooster-Nourish medium (Rooster Bio, Frederick, MD). Cells were counted and cultured as described below

### Cell counting

A Cellometer Vision (Nexcellom, Lawrence, MA) was used to determine total viable cell counts. 20 μl ViaStain AOPI reagent (Nexcelom) was added to an Eppendorf tube containing 20 μl of cells. Once mixed, 20 μl of the solution was added to a Cellometer slide and total cells, live cells, and viability were calculated.

### Cell Culture

Fresh cells were plated in CellBIND^®^ T-225 flasks (Corning, Tewksburry, MA) at a density of 25,000 viable cells/cm^2^ in RoosterNourish™ medium (Rooster Bio, Frederick, MD). Nonadherent cells were removed after the first media change on day 1. Media was then changed every 3-4 days until colonies were ~80-90% confluent (12-14 days). Cells were released with TrypLE (ThermoFisher Sceintific, Waltham, MA). Passaged cells were plated at a density of 3,000 cells/cm^2^ but otherwise followed the same protocol as freshly plated cells.

Generation of MCBs from three donors (DD5, DD6 and DD7) was performed in CellBind^®^ Hyperflasks. Fresh, primary digests were initially plated at 25,000 viable cells/cm^2^ as above. Cells were released with TrypLE and expanded one more passage to form the MCB. The bulk of passage 1 cells were resuspended in cryopreservation medium (CryoStor CS10; BioLife Solutions, Bothell, WA) and stored in the vapor phase of liquid nitrogen.

Cells were passaged up to nine times in a medium composed of DMEM (Cat#10567014, ThermoFisher, USA) and pathogen reduced human platelet lysate (nLivenPR™, COOK Regentec, USA), with and without the addition of ascorbic acid (248 μM; Cat# A2218, Sigma, USA), recombinant basic fibroblast growth factor (10 ng/ml; Cat#233-GMP-025, R&D Systems, USA) and recombinant epidermal growth factor (10 ng/ml; Cat#236-GMP-200, R&D Systems, USA). Cells at 70-80% confluency were harvested and total cell counts were obtained. A portion of the cells was replated at 3000 cells/cm^2^ in triplicate wells of a six-well plate, with media changes every 3-4 days.

### Phenotypic Analysis of MSC via Flow Cytometry

At passages 2, 3 and 4, 1.8 μl of the following single fluorescently-conjugated antibodies or dye, CD3, CD14, CD19, CD31, CD34, CD45, HLA-DR, CD73, CD90, CD105, Stro-1, and 7AAD (Supplemental Table S1), were added to different wells of a 96-well V-bottom plate. 100 μl of MACS (Miltenyi Biotec, Auburn, CA) buffer and 100 μl of cells (200,000 cells) were added to each well containing an antibody. The plate was incubated at 4°C for 30 minutes shielded from light and afterward, the plate was centrifuged for 5 minutes at 300 × g. Cells were washed and resuspended in 200 μl of MACS buffer. An ACEA Biosciences NovoCyte 2060R flow cytometer was used for data collection and data was analyzed using NovoExpress software (Acea Biosciences San Diego, CA).

### Trilineage Differentiation of MSC

MSCs at passages 1 or 3 were seeded in wells of a 12-well plate containing 3 ml Mesencult (Stem Cell Technologies, Vancouver, B.C.) each at 8.0×10^4^, 4.0×10^4^, and 2.0×10^4^ for chondrogenesis, adipogenesis, and osteogenesis differentiation. Wells containing 4.0×10^4^ MSCs in Mesencult medium served as controls. After incubating for 2 hours, Mesencult in the chondrogenesis well was replaced with StemPro chondrogenesis medium (Thermo Fisher Scientific, Waltham, MA). After one day, Mesencult in the adipogenesis and osteogenesis wells was aspirated and replaced with StemPro adipogenesis medium and StemPro osteogenesis medium, respectively. Respective differentiation media were replenished every 3 days as well as Mesencult in control wells. After 14, 12, and 16 days, wells containing chondrocytes, adipocytes, and osteocytes, respectively, were aspirated of media, washed twice with DPBS, fixed with 4% formalin for 30 minutes, washed once with DPBS, and stained. Alcian Blue, which stains chondrocyte proteoglycans blue, in 0.1 N HCl was added to the chondrocyte wells for 30 minutes, the stain was aspirated, the well was washed three times with 0.1 N HCl and neutralized with distilled water, and chondrocytes were visualized under an inverted light microscope (Nikon). Oil Red O, which stains adipocyte fat globules red, was added to the adipocyte well for 15 minutes, the stain was aspirated, the well was washed three times with distilled water, and adipocytes were visualized under an inverted light microscope. 2% Alizarin Red, which stains osteocyte calcium deposits red, was added to the osteocyte well for 3 minutes, the stain was aspirated, the well was washed three times with distilled water, and osteocytes were visualized under an inverted light microscope. All differentiated cells were qualitatively analyzed by visualization of color and morphology.

### Quantitative RT-PCR analysis of gene expression during differentiation

The RNA was isolated from differentiated cells using TRIzol RNA Isolation reagents (Invitrogen, USA) and cDNA was produced using High-Capacity cDNA Reverse Transcription Kit (Applied Biosystems™, USA). The primers (Supplemental Table S2) were obtained from Integrated DNA Technologies Inc. (USA) and were used to measure the gene expressions of the adipogenesis-related proteins lipoprotein lipase (LPL) and fatty acid binding protein 4 (FABP4), osteogenesis-related proteins osteonectin, osteopontin and collagen type-1, and chondrogenesis-related proteins aggrecan, collagen type-2, and SOX9. The real time PCR assays were run on Bio-Rad C-1000 Touch (Bio-Rad, USA) Real-Time PCR System. The data were analyzed relative to the housekeeping gene, GAPDH, then the fold change of gene in differentiated cells was calculated relative to undifferentiated cells.

### Population doubling time

Population doubling time was determined at each passage by using the formula: t*log(2)/log(T_1_/T_0_), where t is the time (hours) between initial plating and cell harvest at 90% confluency, T_1_ is the cell count at harvest and T_0_ is the initial count at seeding.

### CFU-F assays

For freshly digested cells, 5 ml Mesencult, 20 μl Amphotericin B, and 100 μl Gentamycin were added to three wells of a 6-well plate. 2.5×10^5^, 5.0×10^5^, and 7.5×10^5^ cells were added to the first, second, and third wells, respectively. Plates were placed in the incubator until colonies were 90% confluent or up to 12 days. Media was changed every 3-4 days for 14 days. Plates were washed twice with DPBS, and 2 ml methanol was added to each dish for 5 minutes to fix the cells. After 5 minutes, the methanol was decanted, the plate was allowed to air dry and colonies were stained with a 1% crystal violet solution. Colonies containing >50 cells were scored. Passaged cells were assayed similarly except that cells were plated at densities of 32 cells/cm^2^, 65 cells/cm^2^, and 125 cells/cm^2^.

### T cell suppression assays

Suppression of T cell activation was performed according to previously published protocols with minor modifications [26]. Briefly, peripheral blood mononuclear cells were isolated from whole blood (10 ml) by Ficoll (GE, Chicago, IL) separation and resuspended in DPBS. The majority of cells were labeled with carboxyflourescein diacetate succinimidyl ester (CFSE; Sigma, St. Louis, MO) and frozen until used [27]. Passage 2 or 3 MSCs, in some cases pre-stimulated with 100 ng/ml interferon-γ (IFNγ; RnD Systems, Minneapolis, MN) for 18-24 hours, were resuspended in Rooster Nourish (RoosterBio, Frederick, MD) and added to a 96 well flat bottom plate at 4×10^5^, 1×10^5^, 5×10^4^, 2.5×10^4^, 1.5×10^4^, 5×10^3^ cells/well. Rooster Nourish was added to each well until the volume was 200 μl/well. The plate was placed in a 37°C incubator with 10% CO_2_ at 5% humidity for at least two hours to allow MSCs to attach. Cryopreserved PBMCs were quickly thawed and resuspended at a concentration of 4×10^6^ cells/ml in Eagle's minimal essential medium (EMEM; Stem Cell Technologies; supplemented with 10% FBS, 100 μg/ml PenStrep, 2 mM L-glutamine, and 100 μM β-mercaptoethanol). The medium was aspirated from the plates containing MSC and 100 μl of PBMCs were added to all wells containing MSCs as well as wells without MSCs. T cells were stimulated by adding 100 μl of supplemented EMEM with 40 μg/ml phytohemagglutinin (PHA; Sigma-Aldrich, St. Louis, MO) to each well containing MSCs and PBMCs. Control wells containing CSFE-labeled and unlabeled PBMCs alone were also included, half of which were stimulated with PHA and half which were not. The plate was returned to the incubator. After 4 days, PBMCs from each well were removed and labeled with 5 μl CD3-PE and 5 μl of 7AAD before performing flow cytometry

### Statistics

GraphPad Prism version 8 was used for statistical analysis (Student's *t* Test). An P value < 0.05 was considered significant.

## RESULTS

A typical vertebral column (typically T8-L5) before and after removing soft tissues, separating VBs and then fragmenting to sizes of approximately 1.5 cm^3^ is shown in Figure 1. Plastic adherent vBA-MSC possessed a typical spindle-shaped morphology in culture (Figure 1D). Cells from donors were expanded through passage 4 (the initial plating was considered passage 0) and assayed by flow cytometry. vBA-MSC at passages 1-4 expressed very low levels of hematopoietic cell surface markers CD14, CD19, CD34 and CD45 and expressed low to non-existent amounts of human leukocyte antigen DR (HLA-DR) (Figure 2A). The gating strategy and representative flow cytometry dot plots are shown in Supplemental Figure S1. Levels of PECAM1 (CD31)-expressing cells (typically endothelial cells and monocytes) were also low (<7%) at passage 2 (data not shown). Conversely, passaged vBA-MSC were uniformly positive for CD73, CD90 and CD105. Thus, vBA-MSC possess the characteristic MSC surface marker profile [28]. In addition, a variable portion (approximately 20% or lower, depending on the passage number) of the population also expressed the multipotential MSC surface marker Stro-1 [29–32].

**Figure 1.**
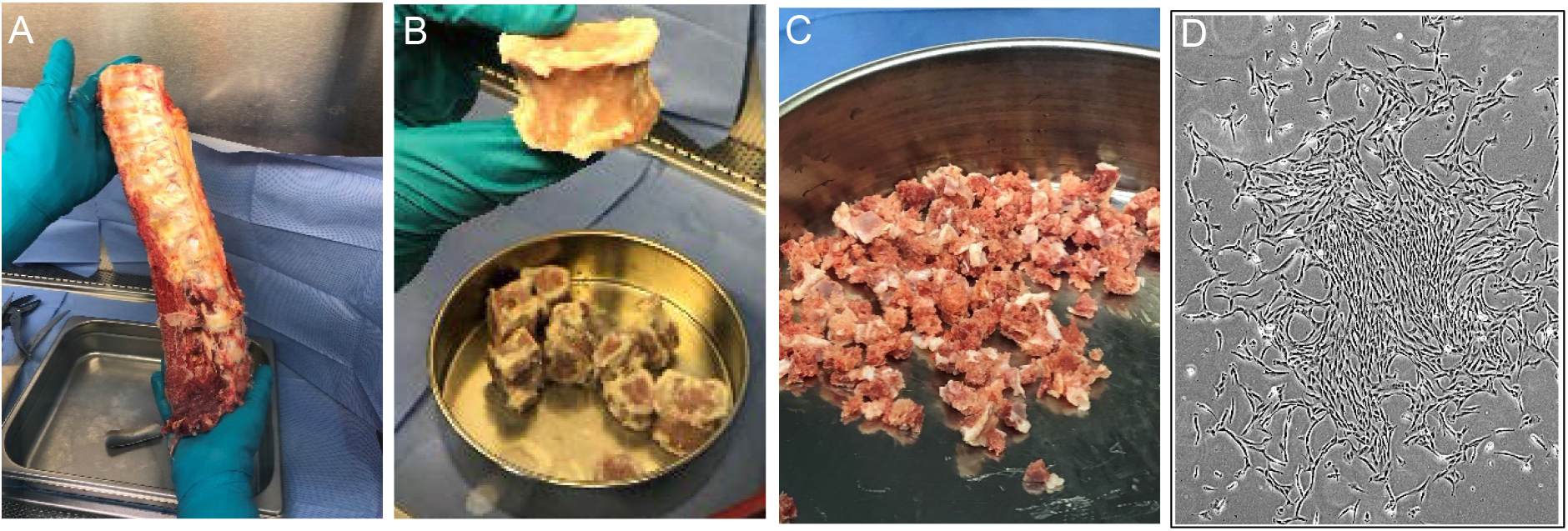
Processing of a typical vertebral column to isolate vBA-MSC. Vertebrae (typically T8-L5) were cleaned of soft tissue (A) before separating vertebral bodies (VBs) and removing disks and remaining soft tissues (B). VBs are ground to approximately 1.5 cm^3^ fragments (C) before enzymatic digestion to release adherent cells. Plastic adherent vBA-MSC form typical spindle shapes in culture (D; passage 2 cells).

**Figure 2.**
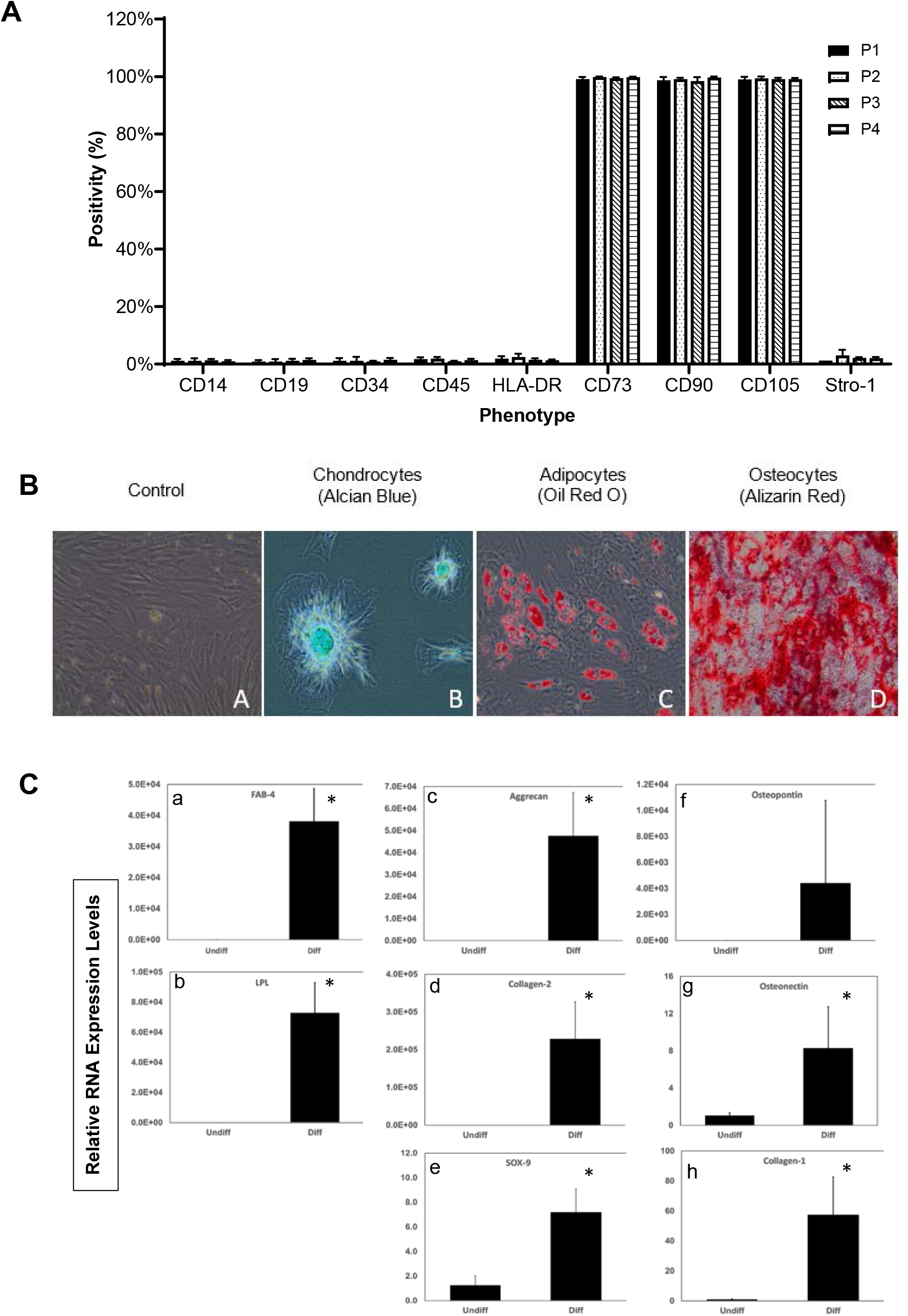
Surface antigen phenotype and trilineage differentiation of vBA-MSC. (A) Passage 1, 2, 3 and 4 vBA-MSC from 3 different donors (DD1, DD2 and DD3; Table 1) were analyzed for surface antigen expression using fluorescently-conjugated antibodies and flow cytometry. The percentage of cells (gated on whole cells using side and forward scatter) after culturing for each passage is shown. (B) Passage 3 vBA-MSC grown in expansion medium (B-A) or induced to undergo either (B-B) chondrogenesis, (B-C) adipogenesis or (B-D) osteogenesis were imaged after staining for chrondocytes (alcian blue), adipocytes (oil red O) or osteocytes (alizarin red), as described in Materials and Methods. Images are representative of results with the 3 different donor-derived vBA-MSC. Magnification for all 20X. (C) Quantitation of differentiation by analysis of (a,b) adipogenic (lipoprotein lipase and fatty acid binding protein-4), (c-e) chondrogenic (aggrecan, collagen 1 and SOX9) and (f-h) osteogenic (osteopontin, osteonectin and collagen 1) RNA markers in undifferentiated (Undiff) and differentiated (Diff) vBA-MSC cultures by quantitative RT-PCR. Relative mRNA levels are shown. *, P<0.05, unpaired Student's t-Test comparing undifferentiated to differentiated cells.

Chondrogenic, adipogenic and osteogenic potentials of passage 3 vBA-MSC were determined for each donor. Each of the vBA-MSC isolates demonstrated the potential to differentiate into chrondrocytes, adipocytes and osteocytes as determined by histology (Figure 2B) and quantitation of RNA transcripts associated with each differentiation pathway (Figure 2C). A portion of both freshly isolated (i.e., never plated) as well as passaged vBA-MSC demonstrated high degrees of clonal proliferation, as determined by colony forming unit-fibroblast (CFU-F) potentials. The average CFU-F frequency in freshly digested VB bone fragments was 0.01±0.004% (mean±standard deviation), which is similar to the frequency of proliferative MSC in whole BM (Figure 3) [7]. The proliferative cells were maintained with cell culture, forming colonies at a frequency of 37±3.4% and 27±1.2% after one and two passages, respectively.

**Figure 3.**
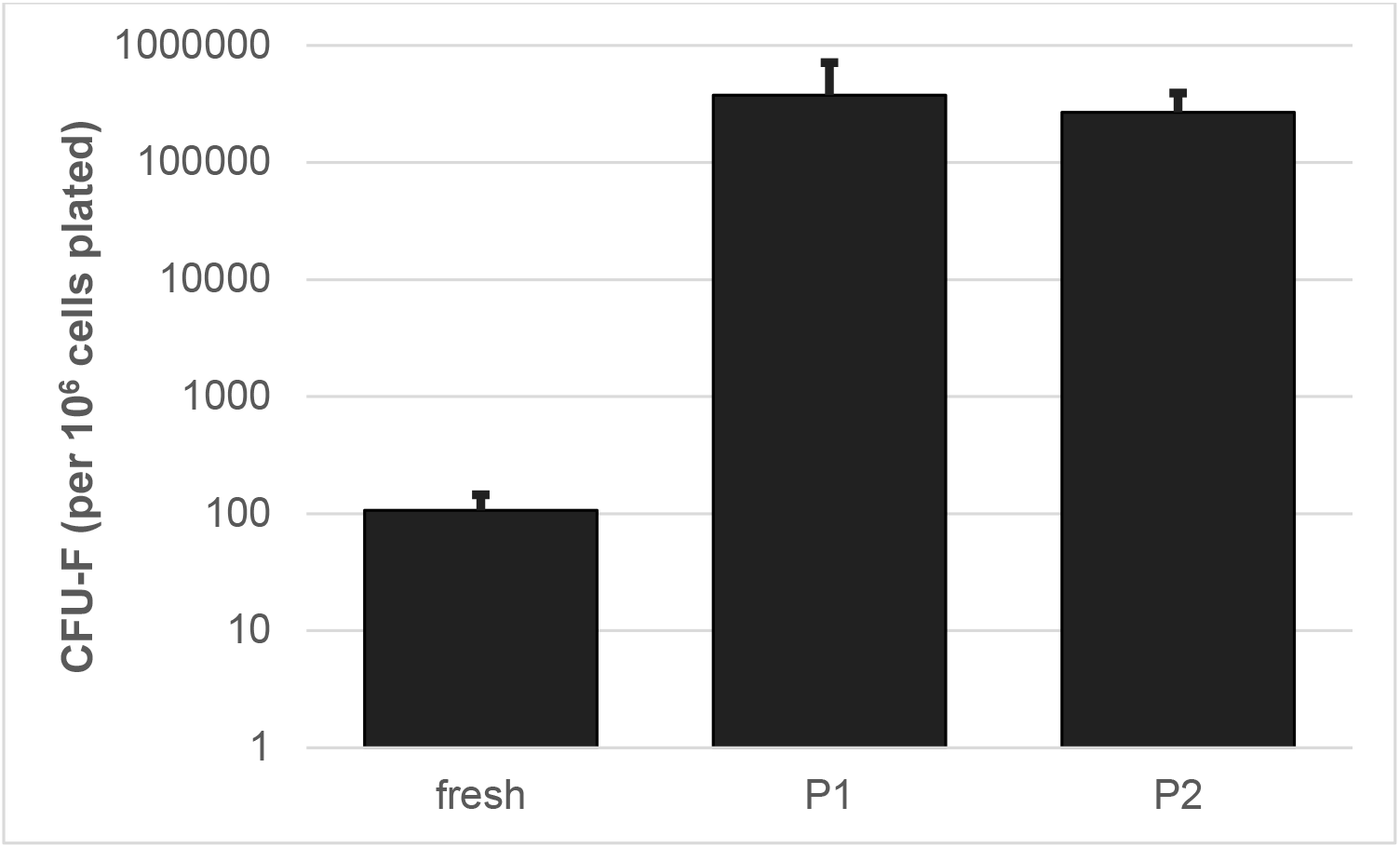
Colony forming unit-fibroblast (CFU-F) potential of vBA-MSC isolated from 3 different donors (DD1, DD2 and DD3; Table 1) and plated immediately after isolation by digestion (fresh) or after 1 or 2 passages (P1 and P2). Either 5×10^5^ (fresh) or 624 (passaged) total cells from each of 3 donors were plated in triplicate wells of a 6 well plate and incubated for 14 days with media changes every 3-4 days.

Suppression of T cell activation is one of the best studied therapeutic properties of MSC, providing the rationale for testing in clinical trials of inflammatory disorders [33, 34]. vBA-MSC from the three different donors dose-dependently suppressed T cells activation with PHA (Figure 4). Maximum suppression at a 1:1 ratio of vBA-MSC to peripheral blood mononuclear cells (PBMC) was 89±7%. A slight but non-significant increase in suppression at all ratios was observed by pre-treating vBA-MSC with IFN-γ for 18-24 hours prior to performing the suppression studies. Treatment with IFN-γ has been shown to stimulate immunosuppressive functions of MSC, with enhanced effects on senescent cells [12]. The insensitivity to priming with concentrations of IFN-γ shown previously to stimulate T cell suppression may indicate that vBA-MSC retain full immunomodulatory capacity during culture expansion.

**Figure 4.**
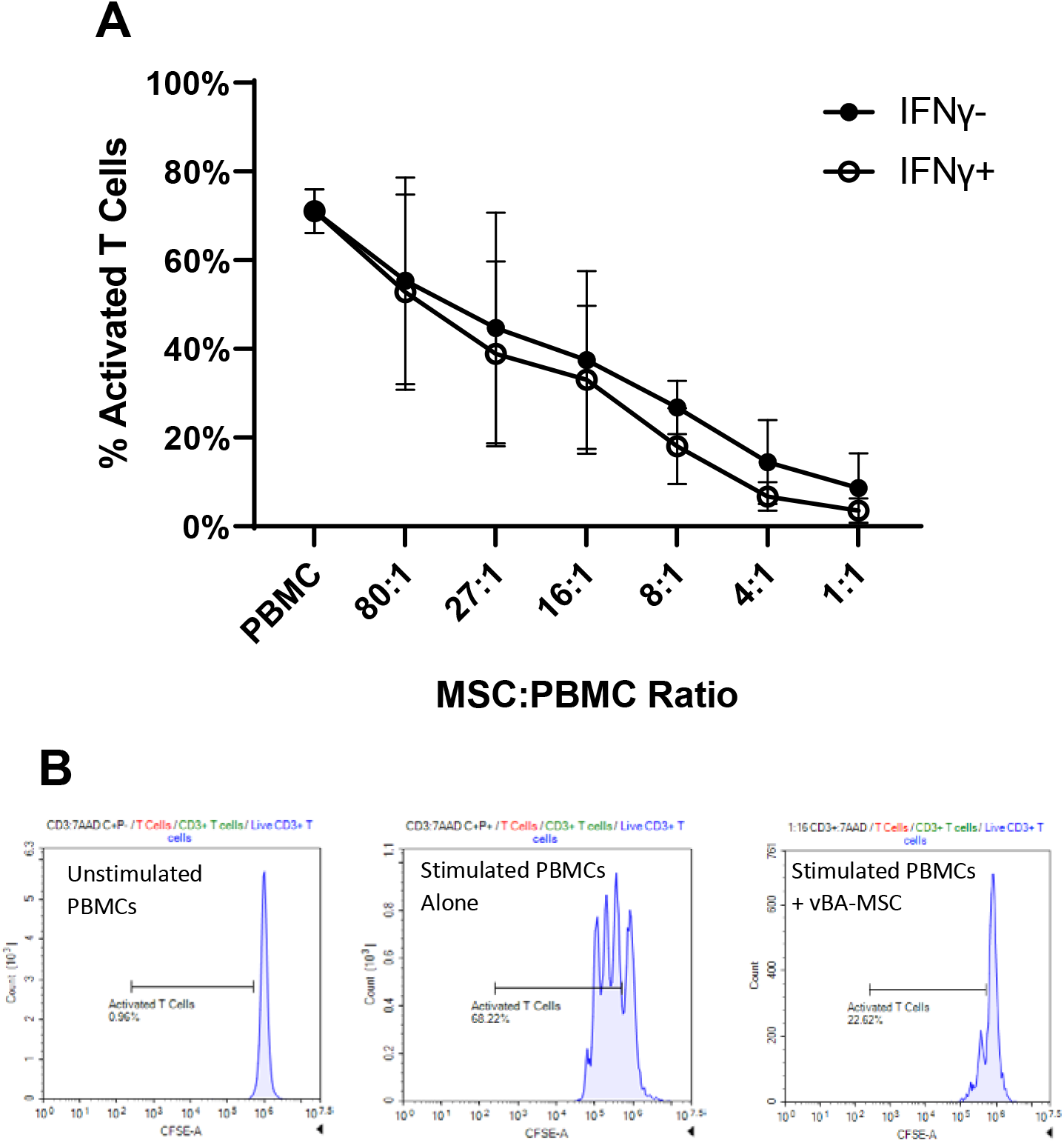
vBA-MSC suppression of T cell activation. (A) Suppression at decreasing ratios of PBMC to vBA-MSC. PBMC isolated from the blood of a single donor were labeled with carboxyfluorescein diacetate succinimidyl ester (CSFE). vBA-MSC were allowed to adhere 2 hours in 96 well plates before washing and adding 4×10^5^ PBMC. In some experiments IFN-γ (100 ng/ml) was added 18-24 hours before adding PBMC. T cells were stimulated for 4 days with PHA. Cells were recovered from the plates and analyzed by flow cytometry after labeling with anti-CD3-PE antibodies. The percentage of activated T cells is plotted. (B) Representative flow plots for PBMCs alone, without and with PHA activation as well as PBMC and MSC after PHA activation are shown. Each data point represents the mean of 3 different experiments with 3 different donors (DD1, DD2 and DD3). Error bars represent the standard deviation. P>0.05 for comparisons at all PBMC:vBA-MSC ratios +/- IFN-γ.

The immunophenotypic profile of plastic adherent vBA-MSC, trilineage differentiation capacity and CFU-F potential as well as immunomodulatory properties confirm the classification of these cells as MSC according to the International Society of Cell and Gene Therapy (ISCT) published guidance [28]. To further establish their equivalency to MSC obtained from BM, a comparison was performed between vBA-MSC and MSC isolated from central BM (Figures 5 and 6). Both commercially available previously expanded live donor BM-MSC (Ex LD BM-MSC), obtained cryopreserved at passage 2, as well as MSC freshly isolated from live donor aspirated BM (LD BM-MSC) were used. In addition, MSC isolated from deceased donor VB BM (DD vBM-MSC) was also included in the comparison. MSC from three donors for each source were expanded to passage 2 and cryopreserved. Upon subsequent thawing, cells were passaged once prior to performing the analyses. MSC from all four sources demonstrated essentially identical immunophenotypic cell surface marker profiles, with very low numbers of cells that expressed CD14, CD19, CD34, CD45 and HLA-DR, and, conversely, nearly all cells expressed CD73, CD90 and CD105 (Figure 5A).

**Figure 5.**
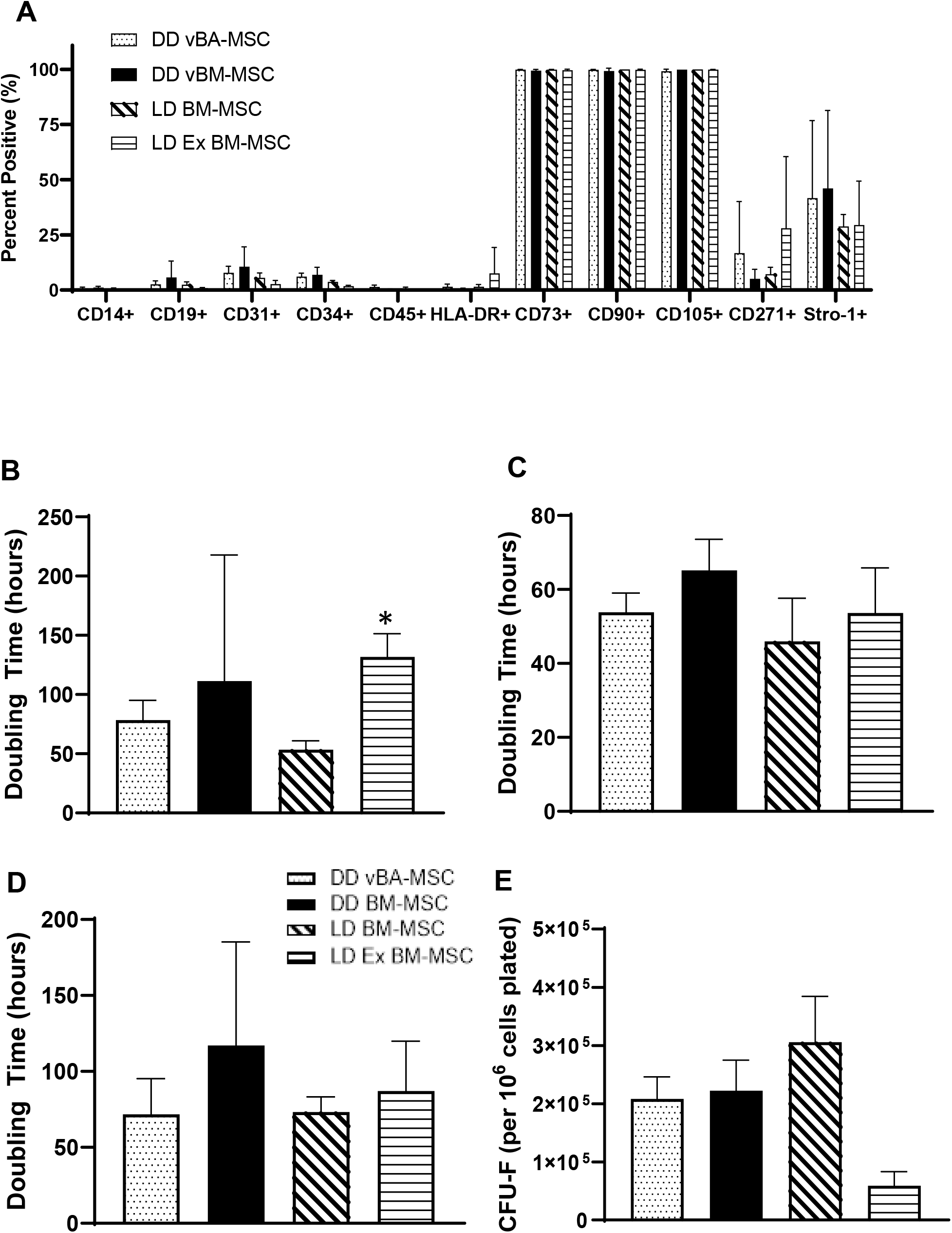
Comparison of vBA-MSC to BM-MSC. (A) Surface marker expression of passage 3 cells was characterized by flow cytometry. The different sources of MSC were: deceased donor vBA-MSC (DD vBA-MSC); deceased donor vertebral body bone marrow-derived MSC (DD BM-MSC); living donor aspirated BM MSC (LD BM-MSC); and living donor aspirated BM MSC obtained from a commercial sources at passage 2 (LD Ex BM-MSC). The percentage of cells within the total population after gating out debris is shown. There were no differences in surface marker expression between cell types. (B-D) Comparison of population doubling times (PDT) from passages 2 to 3 (B), 3 to 4 (C) and 4 to 5 (D). LD Ex BM-MSC grew significantly (*, P<0.05) slower between passage 2 and 3 than either vBA-MSC and LD BM-MSC. No difference in PDT was observed in the subsequent 2 passages. (**E**) CFU-F assays were performed as described in Figure 3 for passaged cells. Formation of CFU-F was significantly lower (*, P<0.05) for passage 2 LD Ex BM-MSC compared to the other three sources of MSC, also at passage 2. Each bar represents the mean ± sd from the 3 donors for each MSC source. The specific donors were: LD BM-MSC (donors LD1, LD2 and LD3); LD Ex BM-MSC (donors LD4, LD5 and LD6); vBM-MSC and vBA-MSC (donors DD1, DD2 and DD3). Donor characteristics are listed in Table 1.

**Figure 6.**
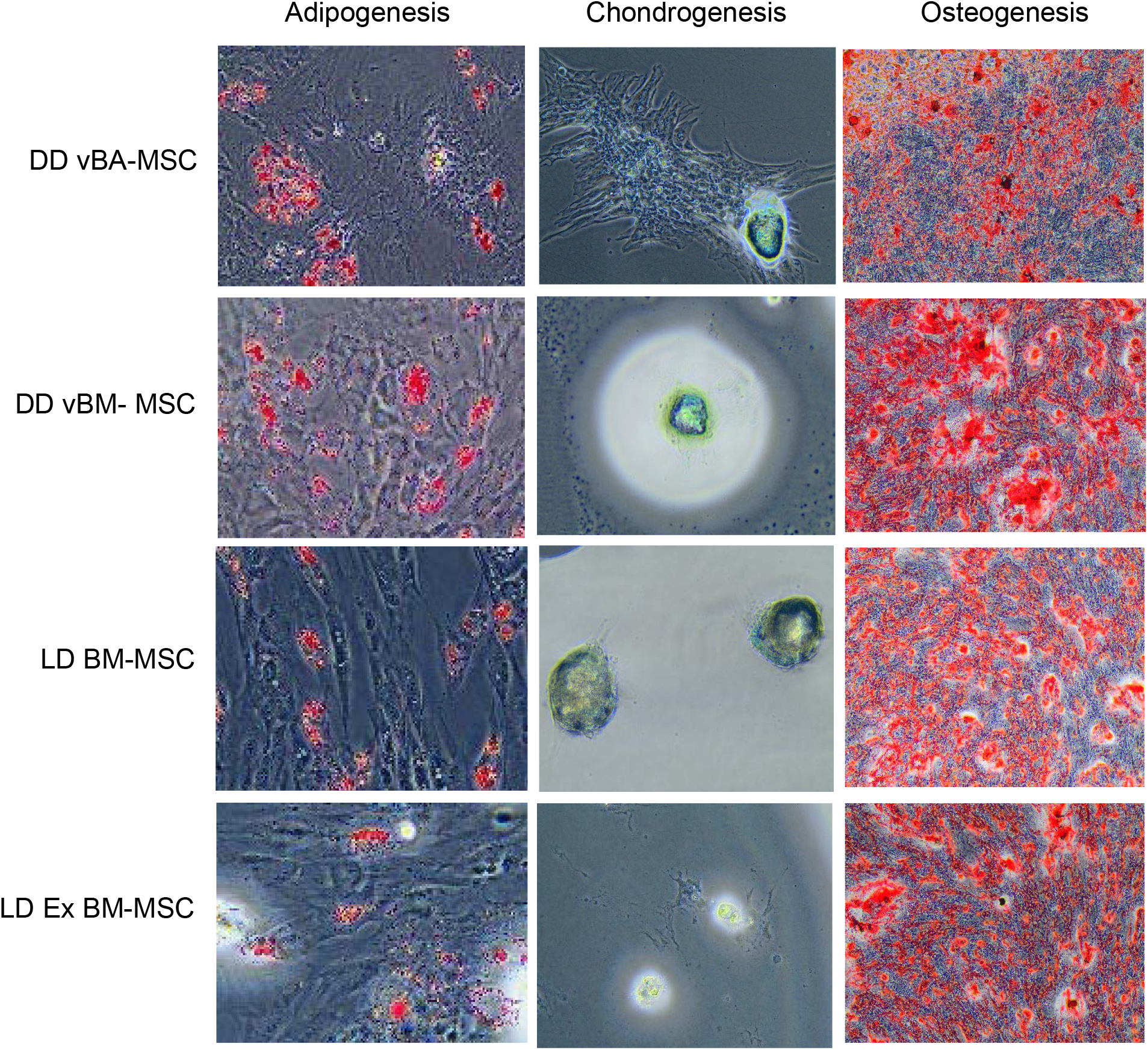
Trilineage differentiation of vBA-MSC and MSC isolated from deceased donor vertebral body BM and BM aspirated from the iliac crests of living donors. Cells were cultured and induced to undergo differentiation for each cell type as described in Figure 2. There were no qualitative differences in either adipogenic, chondrogenic and osteogenic potential of passage 3 cells from any of the four sources. Images are representative from experiments with the 3 different donors for each source of MSC. Magnification is indicated on each image.

MSC from each source grew rapidly in culture through 5 passages (the longest period examined) with no differences in population doubling times (PDTs) at passages 4 and 5 (Figure 5C and D). The Ex LD BM-MSC, which were obtained pre-expanded to passage 2, did exhibit significantly higher PDTs at passage 3 than the other thee MSC populations (Figure 5B). The CFU-F potential of passage 2 Ex LD BM-MSC was also significantly lower than the other MSC populations (Figure 5E). Later passages were not compared for CFU-F potential. Finally, trilineage differentiation potentials were compared and it was found that each MSC population formed adipocytes, chondrocytes and osteocytes *in vitro* at qualitatively the same frequencies (Figure 6).

The potential clinical translational utility of vBA-MSC was assessed by performing a pilot-scale manufacturing run to examine feasibility of banking and expanding large numbers of cells from individual donors. Fragments of VB from 3 different donors were digested to isolate vBA-MSC. The amount (5 g) corresponds to approximately 1/60 of the total VB bone fragment weight (300 g) obtainable from typical donors. Cells enzymatically liberated from VB fragments were plated in a DMEM/human platelet lysate base medium with and without the addition of growth factors and ascorbic acid (see Materials and Methods section). The addition of FGF-2 and EGF was required for optimal growth rate and final yields (S. Thirumala, manuscript in preparation), as demonstrated previously [35]. An MCB at passage 1 from each donor, containing an average of 2.9×10^8^±1.35×10^8^ vBA-MSC, was prepared and the bulk cryopreserved, while the remainder was cultured over multiple passages, tracking total cell yields at each passage (Figure 7A). Passage 1 was considered to be optimal for an MCB, displaying essentially the same surface marker profile and CFU-F potential as later passages (Figures 2 and 3). A single further expansion to passage 2 was enough to produce an WCB containing 5.17×10^9^±4.3×10^9^ vBA-MSC. Based on observed population doublings, two expansions of the entire WCB were sufficient to manufacture over 2×10^12^ (two trillion) cells. The PDT remained nearly constant between passages 2 and 9, without indications of diminishing growth rate at the upper passage number. However, there were differences in PDTs between donors (Figure 7B). Based on the observed PDTs for each donor, starting with a seed stock of 2 million vBA-MSC, it would require 23, 36 and 29 days to manufacture one trillion cells from the three different donors. These times were calculated using 2-dimensional tissue culture flasks and would likely differ in 3-dimensional bioreactors.

**Figure 7.**
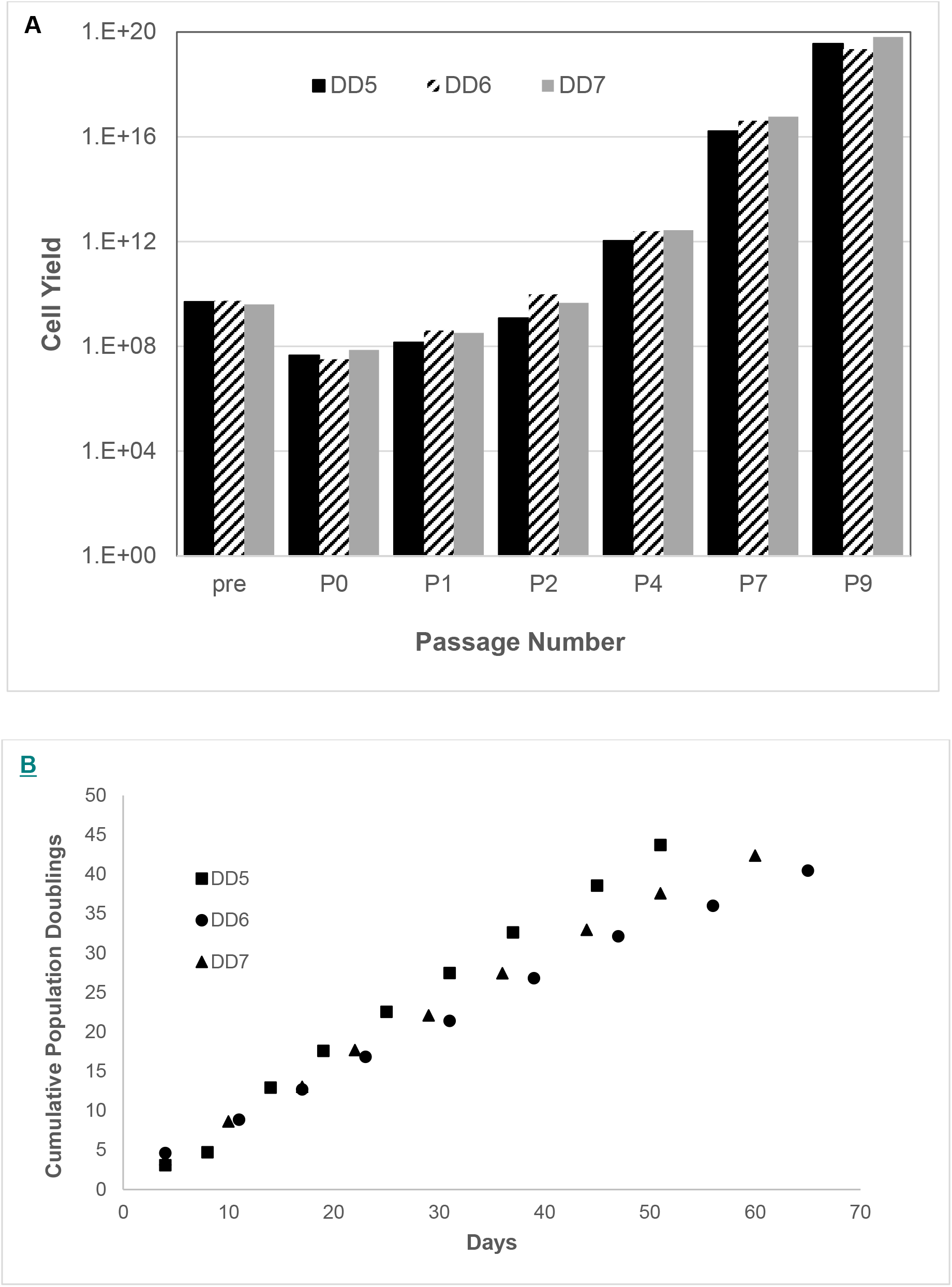
Cumulative population growth of vBA-MSC from 3 different donors. vBA-MSC obtained from digested fragments (5 g) of VBs from 3 donors (DD5, DD6 and DD7) were isolated and expanded to passage 1 to form a master cell bank. A portion of the passage 1 vBA-MSC from each donor was expanded to passage 9. (A) Observed and potential cumulative growth yields at each passage of vBA-MSC from 3 donors. (B) Cumulative vBA-MSC population doublings at passages 0-9. Population doublings (PD) were calculated based on initial numbers of cells plated and the number recovered after each plate reached 80% confluency before replating the cells and was used to determine the theoretical total cell yield after each passage. Theoretical total yields at passages 2-9 were obtained by exponentiating (base 2) the PD calculated for each passage and multiplying by the cumulative cell number from each preceding passage. Each donor vBA-MSC was plated in triplicate for each passage. The coefficient of variation (CV) between cell numbers obtained from each well was <15%.

## DISCUSSION

The transformative potential of MSC to treat a wide variety of medical disorders has been idealized for over a decade; yet, despite many demonstrations of this potential in preclinical and early-stage clinical trials, no MSC-based therapies have achieved success in late-stage, registration (commonly Phase 3 in the U.S.) clinical trials, although a few have received approval for limited indications in relatively small jurisdictions. The reasons for the slow progress in approvals and resulting commercialization of therapeutic MSC despite intense development efforts by multiple entities is certainly multifactorial. In hindsight, it appears that attempts to manufacture MSC at large scale through adopting processes and procedures from the highly successful biopharmaceutical sector might have been a contributory factor [36, 37]. There are many differences between manufacturing products derived from cells versus the cells themselves. Biopharmaceuticals are produced using immortalized cell lines having the ability of nearly unlimited expansion, allowing the generation of large MCBs from a single seed stock. Conversely, the limited availability and expansion potential of MSC requires generating multiple MCB from different donors each year at a disproportionately higher manufacturing costs and regulatory burden [37].

We present here a viable solution to reducing these burdens through the identification and characterization of a large depot of MSC from deceased donor vertebral bones. Based on the analysis presented here, vBA-MSC are phenotypically and functionally equivalent to MSC obtained from central BM. The cells express typical MSC surface markers (CD73, CD90 and CD105) and lack expression of hematopoietic cell surface markers as well as HLA class II proteins (Figures 2 and 5A). Like BM-MSC, vBA-MSC possess the potential to clonally expand and can be induced to undergo trilineage differentiation (Figures 2, 3, 5 and 6). Passaged vBA-MSC are fully fit to suppress T cell activation, demonstrating no difference in activity with prior stimulation by IFN-γ (Figure 4). The differences in PDT and CFU-F of passage 3 (but not later passages) expanded BM-MSC obtained from a commercial source most likely reflects a slower recovery from cryopreservation at passage 2 (Figure 5B). All MSC were grown to passage 2 and cryopreserved in an effort to maintain comparability; however, the commercial source of expanded BM-MSC were likely grown in a different medium and frozen in a different cryopreservation medium. Thus, the cells experienced a lag upon thaw and growth to passage 3 which was not evident in subsequent passages.

Master cell bank sizes averaging 2.9×10^8^ passage 1 vBA-MSC were obtainable from only approximately 1/60^th^ (5 g) of the total digested VB bone fragments recovered from each of 3 donors (Figure 7A). The calculated total yield at passage 4 of vBA-MSC from <u>each donor</u> in this study is over 1×10^14^ (one hundred trillion) cells, equating to over 10^5^ doses at 10×10^6^/kg for an average 70 kg patient. More recent experience, following further optimization of isolation and expansion protocols, suggests an order of magnitude greater yield at each passage is attainable (Ossium internal data). Inevitably, actual total cell yields will be lower due to inefficiencies inherent in large scale manufacturing and requirements for testing; nonetheless, the COG for production of large batches of vBA-MSC from a single donor would likely be much less than for equivalent scales of manufacturing of BM-MSC from multiple donors. The savings in direct manufacturing costs would be in addition to the reduced regulatory burden with using a single donor source for all manufacturing campaigns. The next step in validating the potential cost savings with vBA-MSC will be to perform scaled-up manufacturing runs, which are currently in progress.

We are presently exploring the question of why some populations of MSC are easily dislodged or possibly free floating in the BM, while others remain tightly adhered to the bone/connective tissue matrix and can only be liberated by enzymatic digestion. Determining differences, if they exist, is complicated by the relatively low frequency (<0.01%) of these cells, making them problematic to characterize using common analytical tools, such as flow cytometry, without first expanding in culture, which induces phenotypic and functional alterations [38–46]. One previous report found that freshly isolated enzymatic digests of pelvic region trabecular bone contained 15-fold higher CFU-F than aspirated BM [24]; however, we did not find a similar difference between freshly isolated vBA-MSC and BM-MSC. To better understand dissimilarities, if any, between the populations, we are pursuing single cell RNA sequencing (scRNA-Seq) of vBA-MSC transcriptomes [47, 48]. We are also continuing to characterize the therapeutic potential of vBA-MSC by studying the secretome and extracellular vesicles produced by these cells.

This study was restricted to characterizing vBA-MSC from young, healthy donors between the ages of 15 and 31 years. We also have successfully isolated these cells from older donors (up to 56 years) and demonstrated expansion in culture (unpublished data). However, we intentionally focused on young donors in this report given the literature suggesting higher frequencies and proliferation rates of MSC derived from various tissues obtained from young donors compared to their older counterparts [49–55]. Therefore, in the absence of impacts from environment and disease status, the lowest COG to manufacture vBA-MSC would be from young donors.

In summary, based on the data presented here, the fundamental nature of vBA-MSC does not appear to differ from aspirated BM-MSC; therefore, these cells could potentially be seamlessly substituted for therapeutic applications at a significant savings in manufacturing and regulatory costs. Additionally, other markets requiring large numbers of MSC could also benefit from an abundant source of primary cells. These include tissue engineering and manufacture of products derived from MSC, such as exosomes, as well as biomedical research applications and the emerging applications of cosmeceuticals and bioengineered materials. Each of these markets is expected to grow substantially over the next decades, driving combined demand for MSC in excess of 10 sextillion (1×10^21^) cells annually by 2040 [2]. Future high demand for MSC across all these markets could be entirely met by vBA-MSC obtained from the abundant and steady supply of deceased donor medullary cavity containing bones from the >10,000 organ donors and a further >40,000 tissue donors each year in the U.S. alone [56].

## Acknowlegements

This research was supported by grants from the National Institute of Allergy and Infectious Diseases (AI138334, AI129444) and the National Heart, Lung and Blood Institute (HL142418) to EJW. We thank Jeffrey Gimble MD, PhD at Obatala Sciences, Inc for providing reagents and for reviewing the manuscript. We also thank Nicholas Weinstein for technical assistance.

## Supplemental data

**Supplemental Table S1.** Description of antibodies and dyes used

| Antibody | Fluorophore | Clone | Isotype | Source |
| --- | --- | --- | --- | --- |
| CD3 | PE | UCHT1 | IgG1-PE | BD |
| CD14 | PE | MφP9 | IgG2b-PE | BD |
| CD19 | PE | 4G7 | IgG1-PE | BD |
| CD31 | PE | MBC78.2 | IgG1-PE | BD |
| CD34 | PE | 8G12 | IgG1-PE | BD |
| CD45 | APC | F10-89-4 | IgG2a-APC | Caprico |
| HLA-DR | APC | L243 | IgG2a-APC | Caprico |
| CD73 | PeCy7 | TY/11.8 | IgG1-PeCy7 | Biolegend |
| CD90 | FITC | F15-42-1 | IgG1-FITC | Caprico |
| CD105 | APC | 43A3 | IgG1-APC | Biolegend |
| Stro-1 | APC | STRO-1 | IgM-APC | ThermoFisher |
| 7AAD<sup>1</sup> | -- | -- | -- | Invitrogen |

<sup>1</sup>Abbreviations: 7-AAD, 7-aminoactinomycin; PE, phycoerythrin; APC, allophycocyanin; PeCy7, phycoerythrin-cyanin 7; FITC, fluorescein isothiocyanate.

**Supplemental Table S2.**
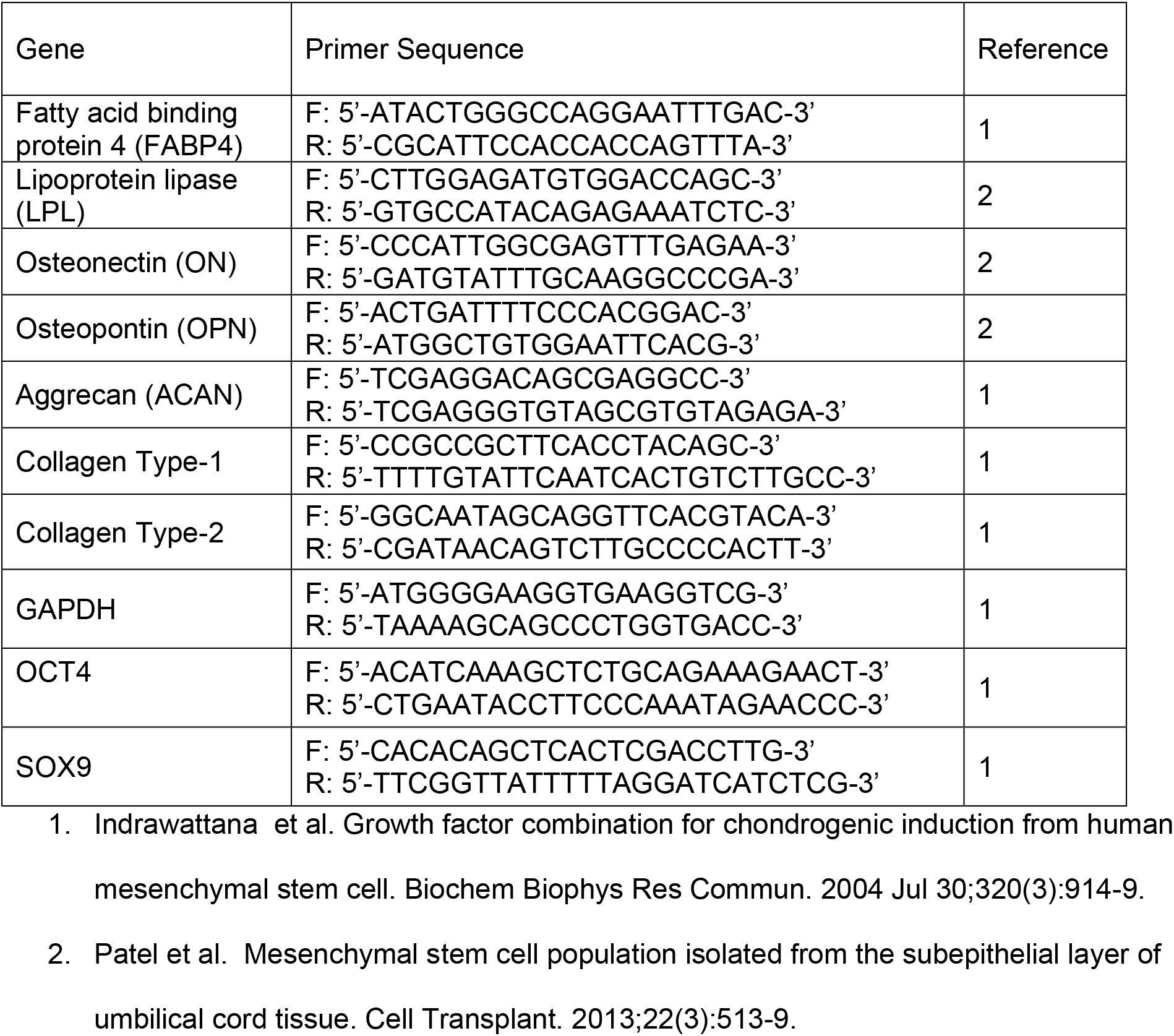
Description of primers used

**Figure S1.**
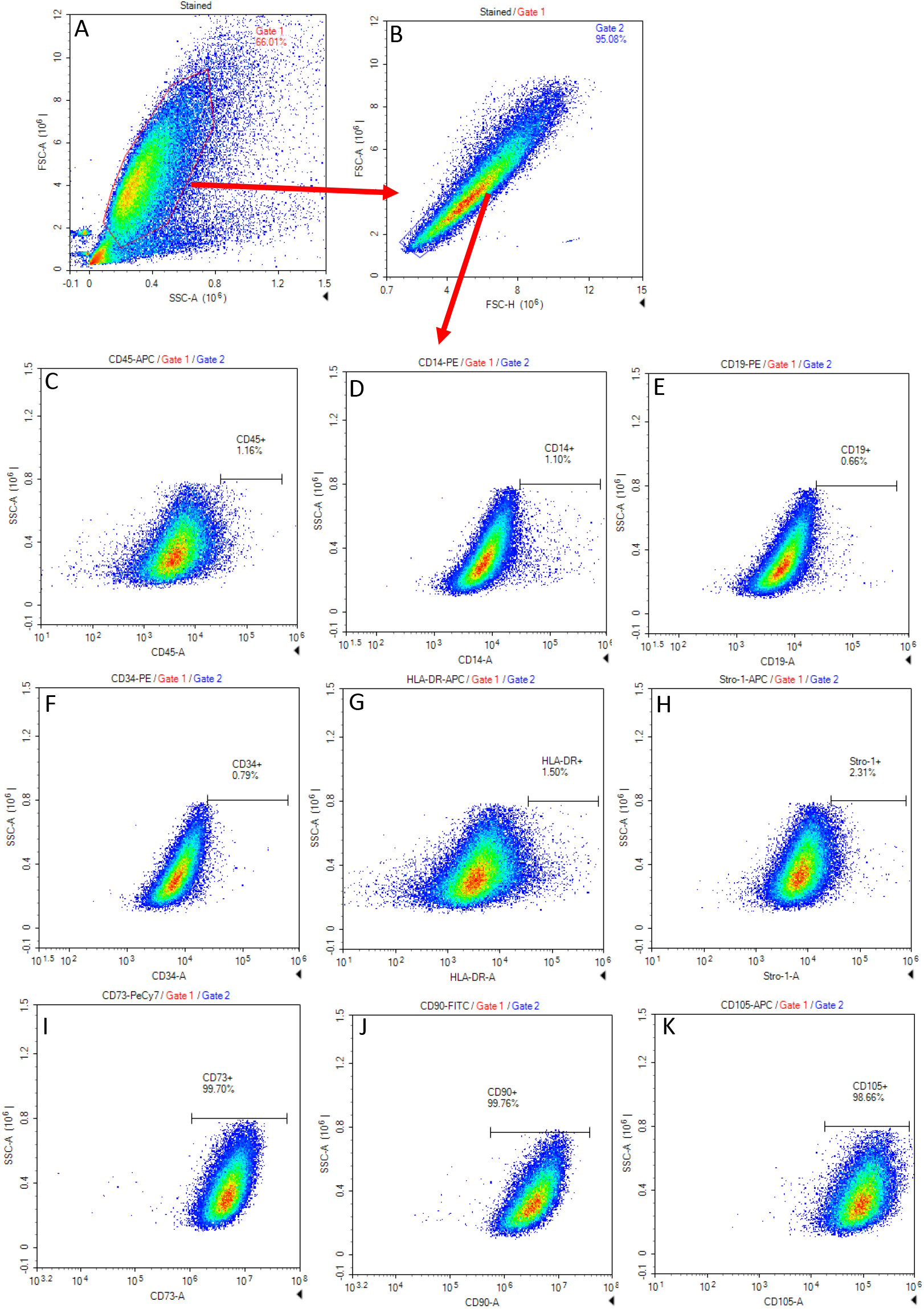
Representative flow cytometry dot plots demonstrating the gating strategy used to identify passaged MSC by size based on forward and side scatter (A) and then single cells by forward scatter area over height (B). Single cells were then analyzed for the indicated cell surface epitopes (H-K) after setting gates based on a negative signal using fluorescently conjugated isotype control antibodies (supplemental table S2).

## Abbreviations

BM: bone marrow
BM-MSC: bone marrow-mesenchymal stem/stromal cell
CFU-F: colony forming unit-fibroblast
COG: cost of goods
DD: deceased donor
LD: live donor
MCB: master cell bank
MSC: mesenchymal stem/stromal cells
OPO: Organ Procurement Organization
PDT: population doubling time
TNC: total nucleated cells
WCB: working cell bank
VB: vertebral bodies
vBA-MSC: vertebral bone-adherent-mesenchymal stem/stromal cells

